# Assessment of behavioral flexibility in mice with conditional deletion of metabotropic glutamate receptor 2 from *Emx1*-lineage neurons

**DOI:** 10.1101/2023.09.15.558004

**Authors:** Doris S. Chang, Mydirah Littlepage-Saunders, Michael J. Hochstein, Christopher J. Matteo, Kidus Amelga, Gregg E. Homanics, Kari A. Johnson

**Affiliations:** Department of Pharmacology, Uniformed Services University of the Health Sciences, Bethesda, Maryland USA; Henry M. Jackson Foundation for the Advancement of Military Medicine, Bethesda, Maryland USA; Neuroscience Graduate Program, Uniformed Services University of the Health Sciences, Bethesda, Maryland USA; Departments of Anesthesiology, Neurobiology, and Pharmacology & Chemical Biology, University of Pittsburgh, School of Medicine, Pittsburgh, Pennsylvania USA

**Keywords:** metabotropic glutamate receptor, habit, behavioral flexibility, alcohol, cocaine

## Abstract

Convergent lines of evidence from animal models suggest that disrupted metabotropic glutamate receptor 2 (mGlu_2_) function promotes high levels of drug consumption for a variety of psychoactive drugs including alcohol, opioids, and psychostimulants. In both rodents and humans, impaired behavioral flexibility prior to first drug use correlates with high levels of drug consumption later in life. Thus, we posited that deletion of mGlu_2_ from brain regions that contribute to behavioral flexibility, including cortical regions, could predispose animals to high levels of drug consumption by impairing behavioral flexibility. To evaluate the role of mGlu_2_ in behavioral flexibility, we generated mice with a floxed *Grm2* allele (*Grm2^f/f^*) and selectively disrupted mGlu_2_ expression in neurons of the *Emx1* lineage (primarily telencephalonic projection neurons) by crossing these mice with an Emx1-IRES-Cre driver line. Behavioral flexibility, including sensitivity to change in either outcome value or action-outcome contingency, was evaluated in adult male and female mice trained to press a lever for a food reinforcer. Contrary to our hypothesis, mGlu_2_ deletion did not facilitate habitual responding assessed by devaluation, contingency degradation, or omission tests. Male *Grm2^f/f^;Emx1-IRES-Cre^+/-^* mice showed modest impairment in reversal learning compared with littermate controls. Finally, we saw a sex-specific effect of mGlu_2_ deletion on response vigor in male mice trained on a random ratio reinforcement schedule. However, we did not find evidence of a general reduction in motivation in a progressive ratio test. These findings suggest that loss of mGlu_2_ from cortical circuitry is unlikely to create a predisposition to inflexible behavior that facilitates excessive drug consumption.

## Introduction

The ability to flexibly update behavioral strategies in response to changing circumstances is critical for adapting to complex environments. Impaired behavioral flexibility, including increased reliance on habitual action strategies and diminished ability to update actions in response to changing rules governing action-outcome relationships, has been frequently observed in animals following exposure to psychoactive drugs (Barker et al., 2020; Corbit et al., 2014a; Everitt and Robbins, 2016; Melugin et al., 2021; Miles et al., 2003; Spear, 2018; Zhukovsky et al., 2019) (but also see Hogarth, 2020; Vandaele et al., 2021). Dysregulation of cortical output, including corticostriatal circuitry, is commonly implicated in behavioral flexibility deficits associated with drug misuse (Barker et al., 2015; Bobadilla et al., 2017; Corbit and Janak, 2016; Gremel and Lovinger, 2017; Izquierdo and Jentsch, 2012; Shields and Gremel, 2020). Understanding mechanisms of circuit dysregulation that contribute to maladaptive behaviors related to drug use will be critical for identifying new treatments that reduce harmful drug consumption.

Metabotropic glutamate receptor 2 (mGlu_2_) is a G_i/o_-coupled G protein-coupled receptor that is primarily expressed presynaptically on glutamatergic terminals (Niswender and Conn, 2010). Activation of mGlu_2_ is known to reduce glutamate transmission in several brain regions involved in maladaptive drug use, including the dorsal striatum, nucleus accumbens, and amygdala (Johnson et al., 2017; Kupferschmidt and Lovinger, 2015; Lovinger and McCool, 1995; Lucas et al., 2013; Robbe et al., 2002). Converging evidence from rodent models suggests that disruption of mGlu_2_ function confers vulnerability to high levels of drug consumption (Jordan and Xi, 2021). Rats lacking mGlu_2_ self-administer high levels of both cocaine and heroin (Gao et al., 2018; Yang et al., 2017), and constitutive *Grm2* knockout mice consume more ethanol and show higher ethanol preference than control mice in a voluntary ethanol drinking task (Zhou et al., 2013). Interestingly, several rat strains that were selectively bred for high levels of ethanol consumption were subsequently found to harbor a premature stop codon in *Grm2* (Wood et al., 2017; Zhou et al., 2013). Pharmacological activation of mGlu_2_ reduces voluntary consumption of psychoactive drugs including alcohol, cocaine, nicotine, and amphetamines (Augier et al., 2016; Crawford et al., 2013; Dhanya et al., 2014; Dhanya et al., 2011; Jin et al., 2010; Johnson and Lovinger, 2016, 2020; Li et al., 2016; Liechti et al., 2007; Sidhpura et al., 2010). In addition, a history of drug exposure can disrupt mGlu_2_ function in wild-type rodents. For example, reduced mGlu_2_ expression in pyramidal neurons of the prefrontal cortex has been observed in alcohol-dependent rats, and restoration of mGlu_2_ expression reduces escalation of alcohol seeking (Meinhardt et al., 2013). In addition, our previous work demonstrated that chronic intermittent ethanol exposure during adolescence reduces mGlu_2_-mediated presynaptic inhibition of glutamatergic transmission in both the dorsolateral and dorsomedial striatum of mice (Johnson et al., 2020a). Collectively, these findings suggest that impaired mGlu_2_ regulation of cortical input to subcortical regions associated with drug misuse could promote drug consumption via effects on cognitive and behavioral processes mediated by these regions, including reward processing, motivation, and behavioral flexibility.

In the studies presented here, we evaluated the impact of loss of mGlu_2_ modulation of cortical output on several types of behavioral flexibility by generating mice with a floxed *Grm2* gene and selectively disrupting mGlu_2_ expression in telencephalonic neurons of the *Emx1* lineage. Based on evidence that a history of drug exposure disrupts mGlu_2_ function in corticostriatal pathways that regulate behavioral flexibility, we predicted that this genetic manipulation would bias mice towards use of habitual action strategies and more generally impair behavioral flexibility. Contrary to our hypothesis, we did not find evidence that loss of cortical mGlu_2_ substantially impairs behavioral flexibility in a variety of tests including satiety-specific devaluation, contingency degradation, omission, and deterministic reversal learning. However, in an operant task that promotes goal-directed behavior, male mice lacking mGlu_2_ show reduced vigor in operant responding, particularly under high-effort conditions. Our findings suggest that impaired behavioral flexibility at baseline is unlikely to contribute to the high levels of drug self-administration and consumption observed in rodents lacking mGlu_2_.

## Results

### Conditional deletion of mGlu_2_ from *Emx1*-lineage neurons

To assess how broad loss of mGlu_2_ regulation of cortical output affects forms of behavioral flexibility that are thought to be associated with high levels of drug consumption, we crossed *Grm2^f/f^* mice with *Grm2^f/f^;Emx1-IRES-Cre^+/-^* mice to generate mice lacking mGlu_2_ expression in neocortex and other telencephalonic structures. Neurons of the *Emx1*-expressing lineage include glutamatergic neurons in the neocortex and hippocampus, as well as a subset of neurons in the amygdala and endopiriform nucleus (Gorski et al., 2002). To confirm specific recombination of *Grm2* in telencephalon vs. other structures, we performed qPCR on samples from primary motor cortex and midline thalamus (a diencephalon region known to express mGlu_2_). Relative expression of *Grm2* was significantly lower in motor cortex samples from *Grm2^f/f^;Emx1-IRES-Cre^+/-^* compared with *Grm2^f/f^;Emx1-IRES-Cre^-/-^* mice (2^-ΔCT^ = 0.017 ± 0.002 for *Grm2^f/f^;Emx1-IRES-Cre^-/-^* and 4.06 x 10^-5^ ± 6.02 x 10^-6^ for *Grm2^f/f^;Emx1-IRES-Cre^+/-^*; p < 0.0001, unpaired t test (**Fig. 1a**) Conversely, there was no difference in *Grm2* expression between genotypes in samples from thalamus (2^-ΔCT^ = 0.011 ± 0.002 for *Grm2^f/f^;Emx1-IRES-Cre^-/-^* and 0.012 ± 0.002 for *Grm2^f/f^;Emx1-IRES-Cre^+/-^*; p = 0.79) (**Fig. 1b**). No deleterious effects of mGlu_2_ deletion on gross anatomy, body condition, or general health were observed; however, we noted that birth rates of male and female mice of each genotype significantly deviated from the expected 25% distribution in each group (male *Grm2^f/f^;Emx1-IRES-Cre^-/-^*: 29.3%; male *Grm2^f/f^;Emx1-IRES-Cre^+/-^*: 19.5%; female *Grm2^f/f^;Emx1-IRES-Cre^-/-^*: 27.8%; female *Grm2^f/f^;Emx1-IRES-Cre^+/-^*: 23.4%; chi-square test: p = 0.0065; n = 522) (**Fig. 1c**). Body weights in early adulthood (postnatal day 70-84) were comparable between genotypes in an age-matched cohort (2-way ANOVA, no effect of genotype: F(1,71) = 0.003, p = 0.95; no sex x genotype interaction: F(1,71) = 0.53, p = 0.47; main effect of sex: F(1,71) = 198.7, p < 0.0001) (**Fig. 1d**).

**Figure 1.**
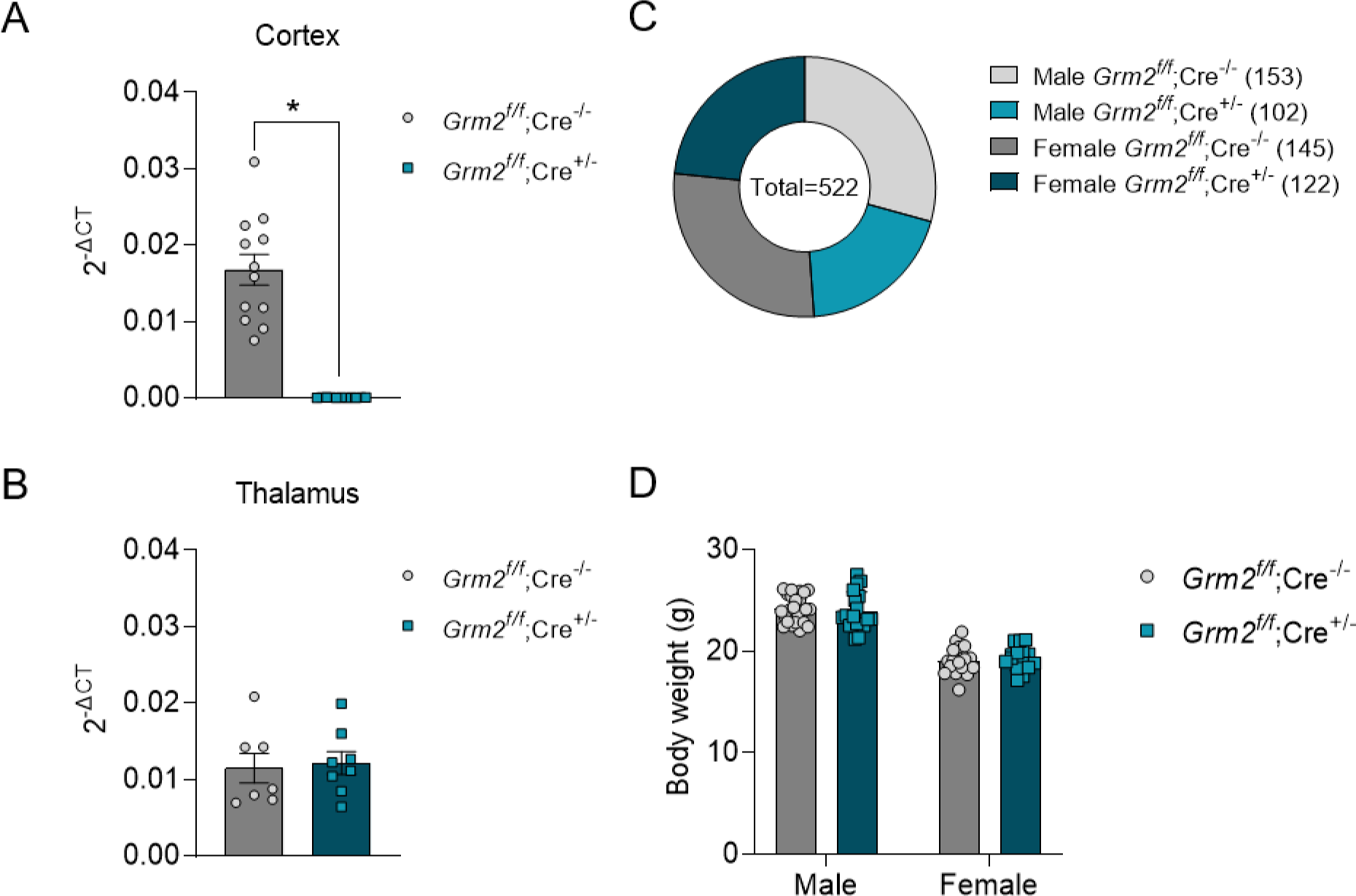
Conditional deletion of mGlu_2_ from *Emx1*-lineage neurons. (A, B) Relative expression of *Grm2* RNA in samples from motor cortex (A; n = 11-12 mice/group) or thalamus (B; n =7-8). Individual data points are averaged from 3 replicates per sample. *p < 0.0001, unpaired t test. (C) Birth rates of *Grm2^f/f^;Emx1-IRES-Cre^-/-^* and *Grm2^f/f^;Emx1-IRES-Cre^+/-^* mice in our colony between June 2020 and March 2023. Numbers represent the total number of mice born and the number born in each group. Chi-square analysis revealed a significant deviation from expected frequencies of 25% for each group (p = 0.0065). (D) Body weights for an age-matched cohort of *Grm2^f/f^;Emx1-IRES-Cre^-/-^* and *Grm2^f/f^;Emx1-IRES-Cre^+/-^* did not differ between genotypes at 70-84 days of age (n = 15-25/group). For panels A, B, and D, data are presented as mean ± SEM with data points from individual mice overlaid.

### Deletion of mGlu_2_ from *Emx1*-lineage cells does not alter performance in tests for habitual behavior or motivation

Our central hypothesis was that disruption of mGlu_2_ function would reduce behavioral flexibility. We first evaluated this by training mice to press a lever for a palatable food reinforcer on a random ratio (RR) schedule, then performing tests that assess sensitivity to changes in outcome value (devaluation) and changes in action-outcome contingency (contingency degradation) (**Fig. 2a**). Because RR training biases towards use of goal-directed actions (Dickinson, 1983; Gremel and Costa, 2013; Lerner, 2020; Yin and Knowlton, 2006), we predicted that training the mice on an RR schedule would provide the best signal window to see a shift towards habitual behavior in mice with mGlu_2_ deletion. Mice escalated rates of lever pressing across RR sessions, with female mice pressing at lower rates than male mice regardless of genotype (3-way ANOVA, main effect of sex: F(1,49) = 82.15, p < 0.0001; main effect of session: F(3.73,182.6) = 409.0, p < 0.0001) (**Fig. 2b,c**). We also found a main effect of genotype (F(1,49) = 7.50, p = 0.0086) and a significant three-way interaction between session, sex, and genotype factors (F(8,392) = 3.897, p = 0.0002). *Post hoc* comparisons revealed that male *Grm2^f/f^;Emx1-IRES-Cre^+/-^* showed lower lever press rates particularly under higher effort conditions (i.e., RR20 responding) (**Fig. 2b**), whereas we did not observe differences in press rates between female *Grm2^f/f^;Emx1-IRES-Cre^-/-^* and *Grm2^f/f^;Emx1-IRES-Cre^+/-^* mice (**Fig. 2c**). We also measured a significantly higher number of head entries into the pellet receptacle in male *Grm2^f/f^;Emx1-IRES-Cre^+/-^* compared with male *Grm2^f/f^;Emx1-IRES-Cre^-/-^*, which was particularly prominent around the transition from continuous reinforcement to RR10 training (2-way RM ANOVA, significant session x genotype interaction: F(8,200) = 4.97, p < 0.0001) (**Fig. 2d**). We did not observe any genotype-related differences in head entries in female mice (**Fig. 2h**). To gain a deeper understanding of how mGlu_2_ deletion affects patterns of operant responding, we performed an analysis of pressing bouts during the final RR20 session (**Fig. 2e-g,i-k**). In male mice, there was no difference in the number of bouts per session (**Fig. 2e**) or the number of presses per bout (**Fig. 2f**), but bout durations were longer in *Grm2^f/f^;Emx1-IRES-Cre^+/-^* mice than in controls (p = 0.004, unpaired t test) (**Fig. 2g**). Calculation of inter-press intervals within bouts demonstrated that male *Grm2^f/f^;Emx1-IRES-Cre^+/-^* mice press significantly slower than controls during bouts (*Grm2^f/f^;Emx1-IRES-Cre^-/-^*: 0.56 ± 0.03 s; *Grm2^f/f^;Emx1-IRES-Cre^+/-^*: 0.71 ± 0.02 s; p = 0.0004, unpaired t test). Consistent with our analysis of press rates across the entire session, we did not observe a genotype difference in bout parameters in female mice (**Fig. 2i-k**).

**Figure 2.**
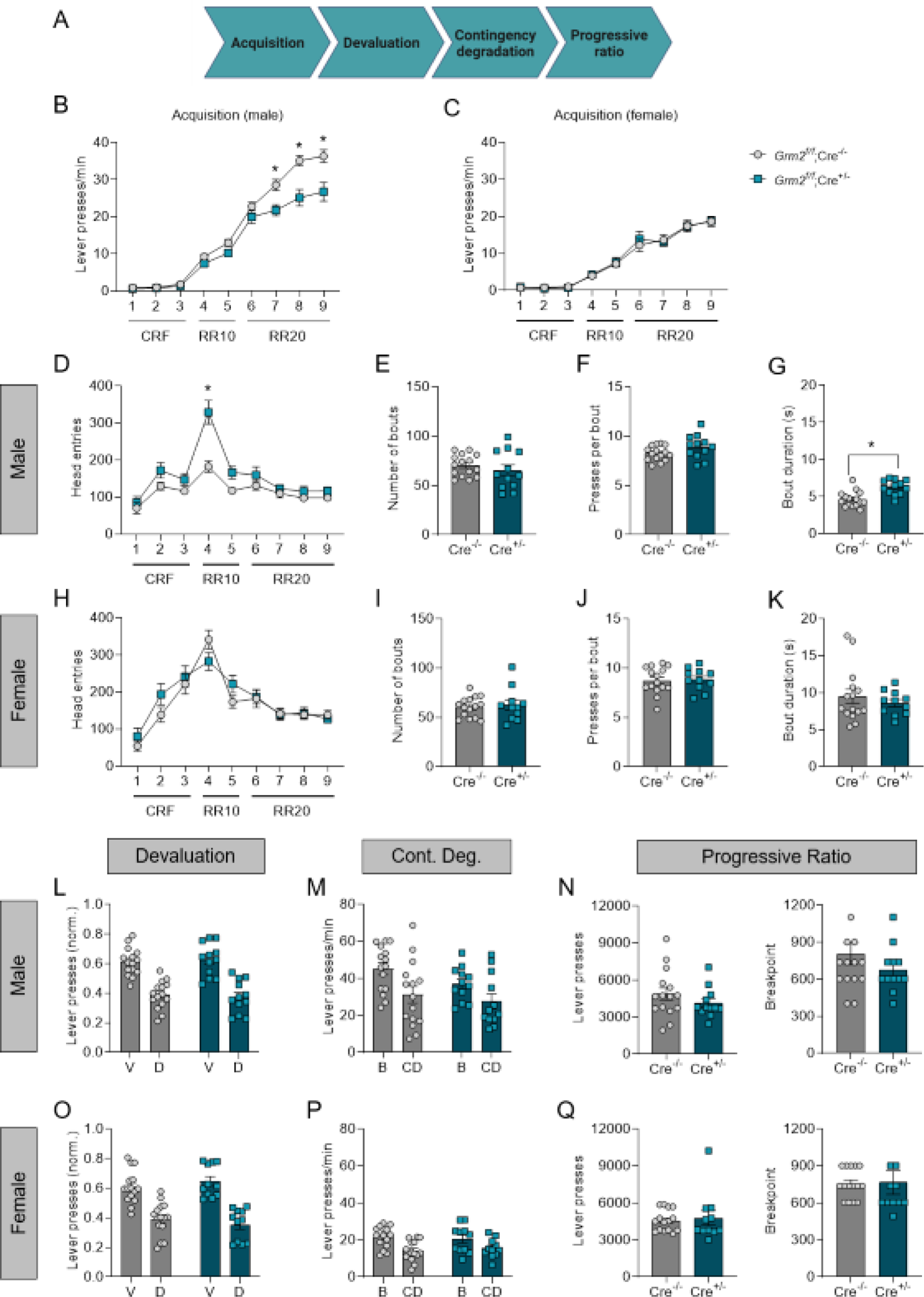
mGlu_2_ deletion does not alter habitual behaviors or motivation following training on a random ratio reinforcement schedule. (A) Experimental timeline included acquisition of lever pressing on a self-paced random ratio (RR) schedule, assessment of habitual action strategies, and progressive ratio testing. (B, C) Lever press rates during acquisition of random ratio lever pressing for a palatable food pellet reinforcer. (B, D-G, L-N) n = 15 *Grm2^f/f^;Emx1-IRES-Cre^-/-^* and 12 *Grm2^f/f^;Emx1-IRES-Cre^+/-^* male mice per group. (C, H-K, O-Q) n = 15 *Grm2^f/f^;Emx1-IRES-Cre^-/-^* and 11 *Grm2^f/f^;Emx1-IRES-Cre^+/-^* female mice per group. (D, H) Head entries per session into the food pellet receptacle during RR acquisition. (E-G, I-K) Analysis of lever pressing patterns during the final RR20 training session. Measures include the number of bouts per session (E, I), the average number of presses in each bout (F, J), and the average duration of each bout (G, K). (L, O) Number of lever presses during valued (V) and devalued (D) 5-minute sessions under extinction conditions. For each mouse, lever presses were normalized by dividing the number of presses made in the valued or devalued session by the total number of presses across both sessions. (M, N) Lever press rates during a baseline (B) RR20 session and during 3 consecutive contingency degradation (CD) sessions. CD data for each mouse were averaged across the 3 sessions. (N, Q) Total number of lever presses (left panels) and breakpoints (right panels) during progressive ratio sessions. Data are presented as mean ± SEM with data points from individual mice overlaid for all bar graphs. *p<0.05 for between-group comparisons of genotype effects (*post hoc* Sidak’s test (B, D) or unpaired t test (G)).

Following training on the RR schedule, mice underwent a satiety-based devaluation test to assess sensitivity to changes in outcome value. Mice pressed the lever less on devalued days than valued days, but deletion of mGlu_2_ did not alter lever pressing in male or female mice (2-way RM ANOVA in male mice: main effect of devaluation, F(1,25) = 36.88, p < 0.0001; no devaluation x genotype interaction, F(1,25) = 0.14, p = 0.71; female mice: main effect of devaluation, F(1,24) = 33.86, p < 0.0001; no devaluation x genotype interaction: F(1,24) = 0.80, p = 0.38) (**Fig. 2l,o**). After re-establishing RR20 responding, mice underwent three contingency degradation sessions (i.e., reinforcer delivery no longer dependent on lever pressing) to assess sensitivity to changes in action-outcome contingency. As expected, mice showed reduced press rates during contingency degradation sessions compared with baseline RR20 response rates; however, mGlu_2_ deletion did not significantly alter sensitivity to contingency degradation (male mice: main effect of contingency degradation, F(1,25) = 23.27, p < 0.0001; no contingency degradation x genotype interaction, F(1,25) = 0.95, p = 0.34); female mice: main effect of contingency degradation, F(1,24) = 29.90, p < 0.0001; no contingency degradation x genotype interaction, F(1,24) = 1.21, p = 0.28) (**Fig. 2m,p**).

After performing devaluation and contingency degradation tests, we also tested mice on a progressive ratio schedule to assess motivation for earning the reinforcer. We did not observe any difference in total lever presses or breakpoints in progressive ratio sessions for either male (**Fig. 2n**) or female mice (**Fig. 2q**).

We trained a separate group of mice to lever press for a food pellet reinforcer on a random interval (RI) schedule. RI schedules have been shown to promote use of habitual action strategies (Dickinson, 1983; Gremel and Costa, 2013; Lerner, 2020; Yin and Knowlton, 2006), and we reasoned that mice with a higher propensity towards habit formation might show a transition to habitual actions earlier in training. For this reason, we included an omission probe to test sensitivity to changes in action-outcome contingency early in training, then proceeded to perform additional tests for habitual behavior after more extensive training on the RI schedule (**Fig. 3a**). Mice escalated lever press rates across sessions, with higher press rates observed in male mice than female mice (3-way ANOVA, main effect of session: F(3.41,190.7) = 107.8, p < 0.0001; main effect of sex: (F1,56) = 35.7, p < 0.0001) (**Fig. 3b,c**). Unlike results obtained from mice trained on an RR schedule, we did not observe a main effect of genotype or any interactions between genotype and sex or session factors in mice trained on the RI schedule. Likewise, we did not find any genotype differences in head entries (**Fig. 3d,h**) or pressing patterns (**Fig. 3e-g,i-k**).

**Figure 3.**
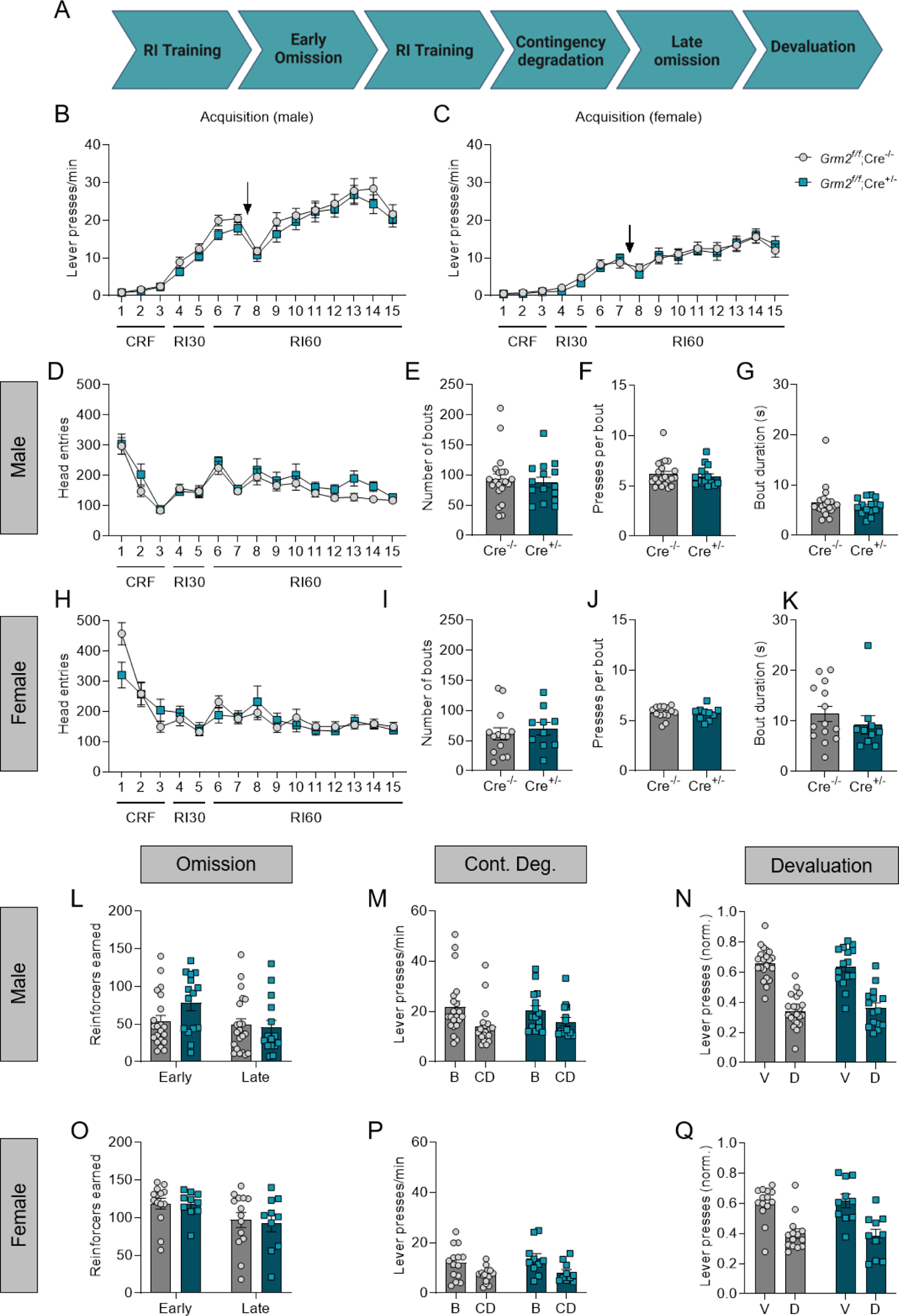
mGlu_2_ deletion does not alter habitual behaviors following training on a random interval reinforcement schedule. (A) Experimental timeline included acquisition of self-paced lever pressing on a random interval (RI) schedule and assessment of habitual actions strategies. (B, C) Lever press rates during acquisition of random interval lever pressing. Arrows indicate the timing of the early omission test. (B, D-G, L-N) n = 21 *Grm2^f/f^;Emx1-IRES-Cre^-/-^* and 15 *Grm2^f/f^;Emx1-IRES-Cre^+/-^* male mice per group. (C, H-K, O-Q) n = 14 *Grm2^f/f^;Emx1-IRES-Cre^-/-^* and 10 *Grm2^f/f^;Emx1-IRES-Cre^+/-^* female mice per group. (D, H) Head entries per session into the food pellet receptacle during RI acquisition. (E-G, I-K) Analysis of lever pressing patterns during the final RI60 training session. Measures include the number of bouts per session (E, I), the average number of presses in each bout (F, J), and the average duration of each bout (G, K). (L, O) Total number of reinforcers earned during 60-minute omission probes early in training and after extensive training (late). (M, N) Lever press rates during a baseline (B) RI60 session and during 2 consecutive contingency degradation (CD) sessions. CD data for each mouse were averaged across the 2 sessions. (N, Q) Number of lever presses during valued (V) and devalued (D) 5-minute sessions under extinction conditions. For each mouse, lever presses were normalized by dividing the number of presses made in the valued or devalued session by the total number of presses across both sessions. Data are presented as mean ± SEM with data points from individual mice overlaid for all bar graphs.

To assess changes in sensitivity to action-outcome contingency, we performed omission probes both early (between the second and third RI60 sessions) and late in training (after ∼10 additional RI60 sessions) (**Fig. 3a**). Consistent with the idea that more extensive training promotes habitual behavior, male mice earned less reinforcers in the late omission test than the early one (2-way RM ANOVA, main effect of session: F(1,34) = 7.21, p = 0.011), but there was no effect of genotype (F(1,34) = 0.97, p = 0.33) and the session x genotype interaction did not reach significance (F(1,34) = 3.87, p = 0.058) (**Fig. 3l**). Female mice also earned less reinforcers in the late omission test than the early one (main effect of session: (F(1,22) = 9.94, p = 0.005), with no effect of genotype (F(1,22) = 0.05, p = 0.83) or session x genotype interaction (F(1,22) = 0.05, p = 0.83) (**Fig. 3o**). In a two-day contingency degradation probe, mice reduced lever press rates relative to a baseline RI60 session (2-way RM ANOVA, male mice, main effect of contingency degradation: (F(1,34) = 48.18, p < 0.0001; female mice, main effect of contingency degradation: (F(1,22,) = 40.54, p < 0.0001). In male mice, we did not find a main effect of genotype (F(1,34) = 0.01, p = 0.92), and a trend towards reduced sensitivity to contingency degradation in male *Grm2^f/f^;Emx1-IRES-Cre^+/-^* did not reach significance (F(1,34) = 3.53, p = 0.069) (**Fig. 3p**). Finally, we performed a devaluation test to assess sensitivity to changes in outcome value. In both male and female mice, we observed significantly less lever pressing on the devalued day relative to the valued day (2-way RM ANOVA, male mice, main effect of devaluation: (F(1,34) = 54.97, p < 0.0001); female mice, main effect of devaluation: (F(1,22) = 16.64, p = 0.0005) (**Fig. 3n,q**). However, no effect of genotype was observed. We note that in this experiment, control mice showed more sensitivity to change in outcome value than is typically reported after extended RI training (Derusso et al., 2010; Dickinson, 1983; Gremel and Costa, 2013).

### Pharmacological activation of mGlu_2/3_ does not alter performance in tests for habitual behavior

To avoid confounding factors such as compensation that can occur when gene expression is disrupted from early development, we also assessed the ability of the mGlu_2/3_ agonist LY379268 to alter use of habitual action strategies when administered acutely before habit probes. This experiment was performed on a smaller number of animals and was therefore analyzed as a mixed male and female cohort. We selected a 1.0 mg/kg dose of LY379268, which was shown to modestly reduce FR10 responding (by ∼10-15% of baseline) in our previous studies (Johnson et al., 2020b) and does not inhibit locomotor activity. We trained adult male and female C57BL/6J mice on a RI schedule without drug treatment (**Fig. S2a**), then assessed RI60 responding and sensitivity to changes in outcome value (devaluation) or action-outcome contingency (omission) following vehicle or LY379268 administration. Unlike our previous report of LY379268 effects on FR10 responding, LY379268 did not significantly change press rates (**Fig. S2b**), the number of press bouts per session (**Fig. S2c**), or bout duration (**Fig. S2e**). We did observe a larger number of presses per bout in LY379268-treated mice relative to vehicle-treated mice (p = 0.022, unpaired t test; **Fig. S2d**), but there was not a significant difference in within-bout inter-press intervals between groups (0.78 ± 0.09 s for vehicle-treated mice, 0.81 ± 0.11 s for LY379268-treated mice, p = 0.81). LY379268 did not alter sensitivity to changes in outcome value following RI training (main effect of devaluation (F(1,17) = 31.05, p < 0.0001; no devaluation x drug interaction (F(1,17) = 0.02, p = 0.90) (**Fig. S2f**). Likewise, LY379268 did not appear to alter sensitivity to changes in action-outcome contingency, as mice given vehicle or LY379268 prior to an omission probe earned a similar number of reinforcers (p = 0.69, unpaired t test) (**Fig. S2g**). Because we did not perform a dose-response experiment, the possibility remains that a higher dose of LY379268 could modulate RI60 responding or use of habitual action strategies.

### Deletion of mGlu_2_ from *Emx1*-lineage cells produces a modest impairment of reversal learning in male mice

In addition to promoting habitual action strategies, prior experience with drugs such as alcohol or cocaine has been shown to disrupt updating of actions in response to rule changes, for example, in reversal learning or attentional set shifting tasks (Calu et al., 2007; Crews et al., 2019; Dannenhoffer et al., 2021; Demer et al., 1989; Gould et al., 2012; Izquierdo and Jentsch, 2012; Kantak, 2020; Kromrey et al., 2015; Porter et al., 2011; Shnitko et al., 2020; Zhukovsky et al., 2019), and poorer baseline performance in cognitive flexibility tasks has been correlated with later risk for high levels of drug consumption (Grant et al., 2021; Melugin et al., 2021; Shnitko et al., 2019). We assessed the effect of disrupting mGlu_2_ function in *Emx1*-lineage neurons in a deterministic reversal learning paradigm that requires the mouse to discriminate between an active and inactive lever based on the lever’s position in the chamber (acquisition phase), then update its lever choice after a reversal of the lever-outcome contingency. Three-way ANOVA revealed a main effect of sex (F(1,68) = 7.751, p = 0.0069) and session (F(4.08,277.7) = 180.8, p < 0.0001), demonstrating that the mice improved performance across sessions during the acquisition phase (**Fig. 4a,b**). We did not observe an effect of genotype during the acquisition phase. During the reversal phase, we observed a modest impairment of reversal learning in male *Grm2^f/f^;Emx1-IRES-Cre^+/-^* mice relative to control mice (2-way RM ANOVA, genotype x session interaction: F(14, 462) = 2.46, p = 0.0023) (**Fig. 4c**). We did not find evidence of changes in reversal learning rates in female mice (2-way RM ANOVA, genotype x session interaction: F(14,364) = 0.76, p = 0.72) (**Fig. 4d**). During the reversal phase, there was no effect of genotype or session on omitted trials in male mice (**Fig. 4e**). Female mice slightly increased the number of omitted trials across the reversal phase (2-way RM ANOVA, main effect of session: F(4.9, 127.7) = 2.69, p = 0.025) (**Fig. 4f**).

**Figure 4.**
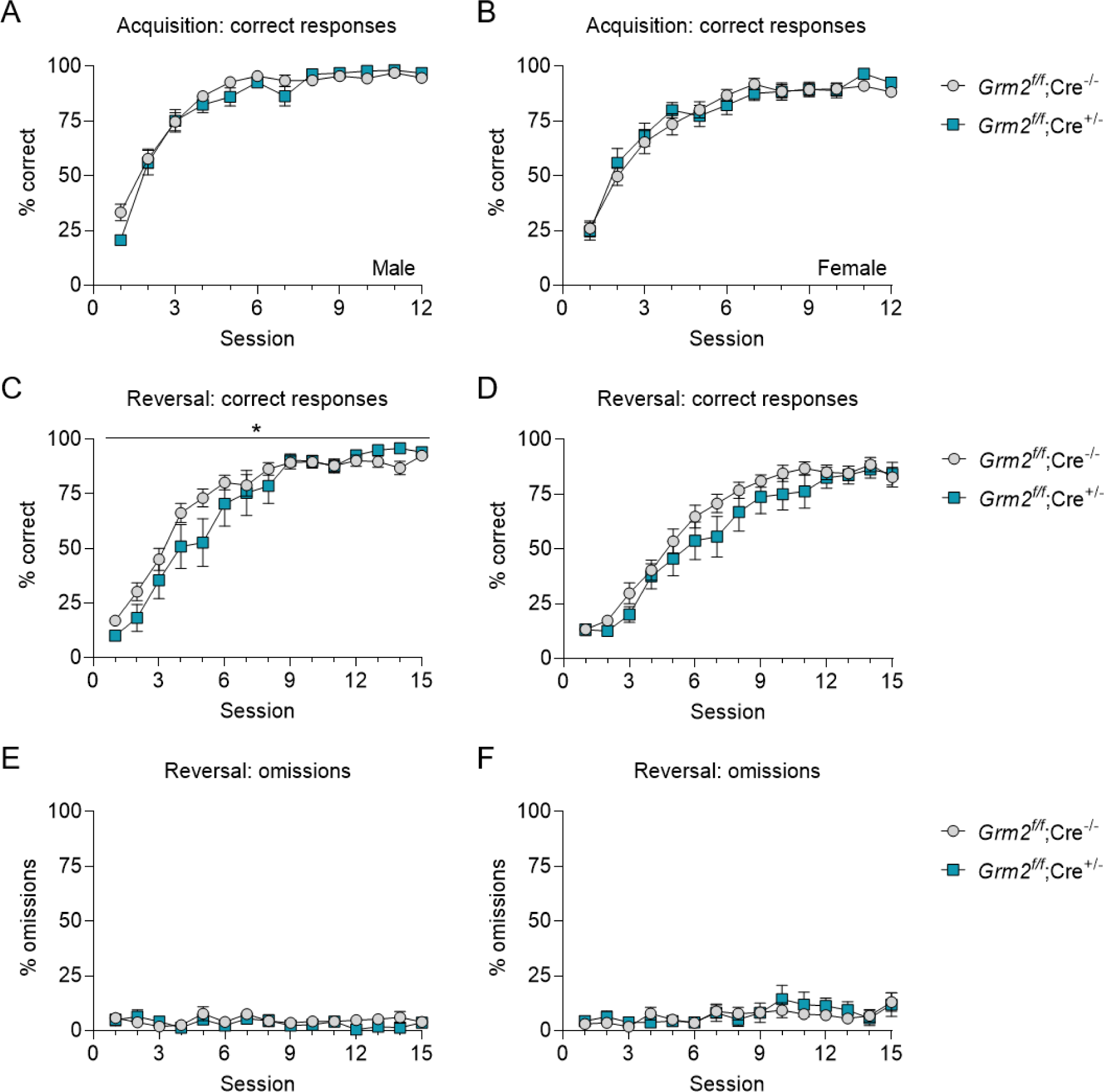
mGlu_2_ deletion produces a modest impairment of reversal learning in male mice. (A, B) Percentage of correct trials during acquisition of a trial-based two-lever place discrimination task. (A, C, E) n = 21 *Grm2^f/f^;Emx1-IRES-Cre^-/-^* and 11 *Grm2^f/f^;Emx1-IRES-Cre^+/-^* male mice per group. (B, D, F) n = 24 (*Grm2^f/f^;Emx1-IRES-Cre^-/-^*) and 13 (*Grm2^f/f^;Emx1-IRES-Cre^+/-^*) female mice per group. (C-F) Percentage of correct trials (C, D) and omitted trials (E, F) after reversal of the active lever. (C) *Significant genotype x session interaction (two-way RM ANOVA, F(14,462) = 2.46, p = 0.0023. Data are presented as mean ± SEM.

### Deletion of mGlu_2_ from *Emx1*-lineage cells does not alter locomotor activity in a novel environment

Our finding that male *Grm2^f/f^;Emx1-IRES-Cre^+/-^* mice show lower rates of lever pressing during acquisition of RR responding for food could be explained by a general decrease in activity levels. To assess this, we measured locomotion in a novel open field apparatus. We found no evidence of reduced locomotor activity in either sex (2-way ANOVA: no effect of genotype, F(1,32) = 0.1, p = 0.76; no sex x genotype interaction, F(1,32) = 0.73, p = 0.40) (**Fig. 5a**). We also measured time spent in the center of the open field and found a main effect of sex (F(1,32) = 4.29, p = 0.047), but no effect of genotype (F(1,32) = 2.84, p = 0.10) or sex x genotype interaction (F(1,32) = 0.25, p = 0.62) (**Fig. 5b**).

**Figure 5.**
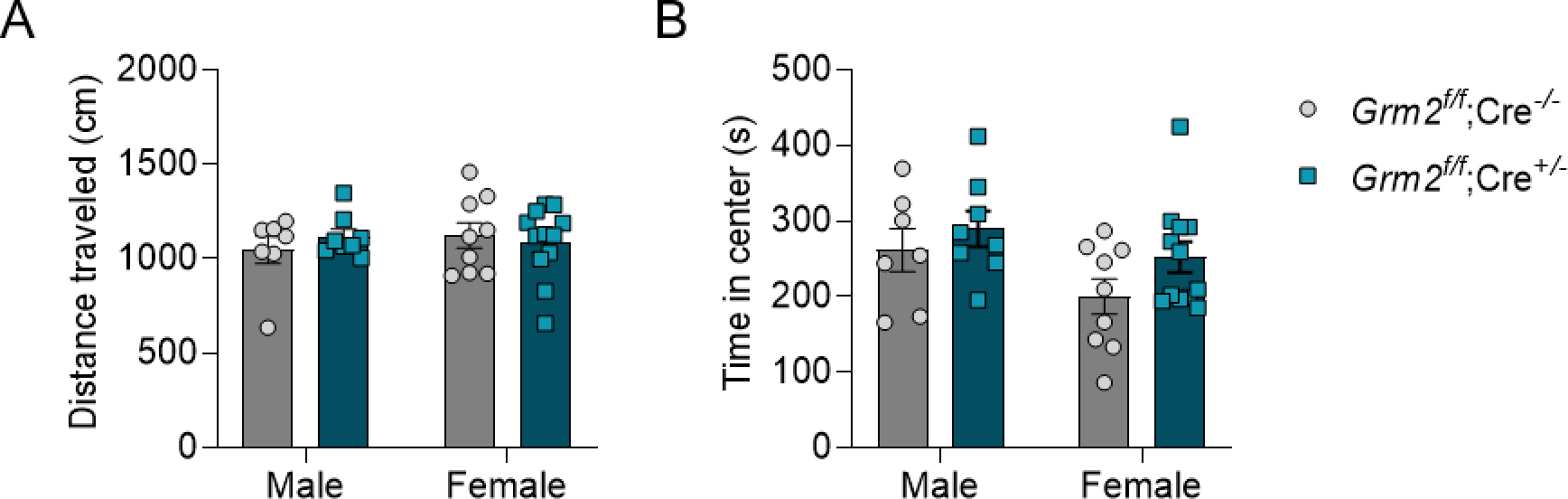
mGlu_2_ deletion does not alter locomotor activity in a novel environment. (A) Distance traveled during exploration of a novel open field apparatus. (B) Time spent in the center of the open field apparatus. n = 7 male *Grm2^f/f^;Emx1-IRES-Cre^-/-^*, 8 male *Grm2^f/f^;Emx1-IRES-Cre^+/-^*, 9 female *Grm2^f/f^;Emx1-IRES-Cre^-/-^*, and 12 female *Grm2^f/f^;Emx1-IRES-Cre^+/-^* mice per group. Data are presented as mean ± SEM with data points from individual mice overlaid.

## Discussion

Because mGlu_2_ deletion predisposes animals to high levels of drug consumption (Jordan and Xi, 2021), and low levels of cognitive flexibility predict high levels of drug consumption (Grant et al., 2021; Melugin et al., 2021; Shnitko et al., 2019), we predicted that loss of mGlu_2_ would impair baseline measures of behavioral flexibility in drug-naïve mice. Our experiments did not demonstrate notable differences in sensitivity to outcome devaluation or adaptation to changes in action-outcome contingencies in mice lacking mGlu_2_ in telencephalon-derived neurons (**Fig. 2,3**). We did observe an impairment in a reversal learning task in male *Grm2^f/f^;Emx1-IRES-Cre^+/-^* compared to controls (**Fig. 4c**), but the effect was modest. Because our experimental design assessed behavioral flexibility in the context of a natural reinforcer (i.e., palatable food), it remains possible that impaired mGlu_2_ function could facilitate transitions to inflexible self-administration of specific psychoactive drugs despite the lack of impairment in drug-naïve mice.

The most prominent effect of mGlu_2_ deletion observed in our experiments was reduced response vigor (both across the session and within lever press bouts) in male mice under high-effort RR conditions (**Fig. 2b**). Due to the paucity of research on sex differences in mGlu_2_-mediated synaptic modulation, it is unclear why mGlu_2_ deletion selectively reduced response vigor in male mice. Reduced response vigor could suggest a generalized impairment of motivation to obtain the food reinforcer. However, we did not observe any difference in breakpoints between *Grm2^f/f^;Emx1-IRES-Cre^-/-^* and *Grm2^f/f^;Emx1-IRES-Cre^+/-^* mice in a progressive ratio test that was performed in the same mice (**Fig. 2n**), suggesting that mGlu_2_ has a specific role in modulating response vigor rather than general motivation under high-effort conditions. Effects of mGlu_2_ deletion on general activity levels are unlikely to account for reduced response vigor in this model, as we did not observe any effect of mGlu_2_ deletion on locomotor activity in a novel open field apparatus. Importantly, our finding that loss of mGlu_2_ reduces vigor when responding for palatable food argues against a generalized enhancement of consummatory behavior that could contribute to increased volitional drug consumption in rodents lacking mGlu_2_.

A limitation of these studies is the relatively broad deletion of mGlu_2_ from telencephalon-lineage neurons using the Emx1-IRES-Cre driver line. We decided to broadly disrupt cortical mGlu_2_ expression based on previous findings that alcohol exposure disrupts mGlu_2_ regulation of glutamatergic transmission throughout the striatal complex (Johnson et al., 2020a; Meinhardt et al., 2013), including dorsal and ventral striatal regions that receive input from many distinct cortical areas (Hunnicutt et al., 2016). It remains possible that more specific deletion of mGlu_2_ in cortical inputs to the dorsomedial, dorsolateral, or ventral striatum could selectively disrupt distinct aspects of behavioral flexibility. For example, disinhibition of cortical inputs to the dorsolateral striatum has the potential to reduce sensitivity to changes in outcome value (Barker et al., 2015; Corbit and Janak, 2016; Gremel and Lovinger, 2017). In rats, habitual alcohol self-administration depends on dorsolateral striatum function, and blocking AMPA receptor-mediated glutamatergic transmission in the dorsolateral striatum reduces habitual control over alcohol consumption (Corbit et al., 2014b).

Consideration of subcortical circuit mechanisms by which mGlu_2_ modulates the rewarding and reinforcing effects of psychoactive drugs will be necessary to understand how impaired mGlu_2_ function contributes to excessive drug intake. Activation of mGlu_2_ modulates striatal dopamine release through actions on subcortical circuitry (Johnson et al., 2017; Johnson et al., 2020b; Littlepage-Saunders et al., 2023). Dopamine signaling in the dorsolateral striatum contributes to habitual responding for natural reinforcers (Lerner, 2020) and for drugs including alcohol, cocaine, and heroin (Belin et al., 2009; Corbit and Janak, 2016; Corbit et al., 2014b; Hodebourg et al., 2019; Willuhn et al., 2012), whereas dopamine transmission in the dorsomedial striatum supports reversal learning (Clarke et al., 2011; Grospe et al., 2018; van der Merwe et al., 2023). Thus, it is possible that deletion of mGlu_2_ from specific subcortical circuits that regulate striatal dopamine transmission could promote inflexible behaviors that facilitate excessive drug intake. Global loss of mGlu_2_ increases basal extracellular dopamine levels in the nucleus accumbens and produces complex effects on dopamine and glutamate responses to psychoactive drugs such as cocaine and heroin (Gao et al., 2018; Yang et al., 2017). Thus, loss of mGlu_2_ could affect the rewarding or reinforcing properties of psychoactive drugs independent of dysregulated cortical circuitry. Improving understanding of the mechanisms (at the levels of both circuits and behavior) by which disrupted mGlu_2_ function predisposes animals to high levels of drug intake could help to identify specific contexts in which pharmacological treatments targeting mGlu_2_ are likely to be most effective.

## Materials and Methods

### Animals

Male and female C57BL/6J and Emx1-IRES-Cre mice were purchased from the Jackson Laboratory (strains 000664 and 005628; Bar Harbor, ME). Breeding colonies of *Grm2flox* and Emx1-IRES-Cre mice were maintained at Charles River Laboratories (Wilmington, MA). Experimental mice were imported to the vivarium at Uniformed Services University of the Health Sciences at 5-10 weeks of age. Upon arrival, mice were group housed (2-5 per cage) in the USU Department of Laboratory Animal Resources rodent vivarium on a 12-hour light/dark cycle (lights on at 0600). Mice were housed on ventilated racks in a temperature-controlled room with *ad libitum* access to food and water, except during testing requiring food restriction. All mice were 2.5-5 months old at onset of behavioral testing. Studies were carried out in accordance with the National Institutes of Health guide for the care and use of laboratory animals and were approved by the Uniformed Services University or University of Pittsburgh Institutional Animal Care and Use Committee. All behavioral testing was conducted during the lights on period.

### Generation of *Grm2^f/f^* mice

CRISPR/Cas9 based mutagenesis was used to create mice with an 1843 bp deletion in *Grm2* and mice with a floxed *Grm2* gene, both on an inbred C57BL/6J genetic background. Details of CRISPR/Cas9 mutagenesis including gRNAs, embryo manipulation, genotyping, off target analysis and creation/characterization of the *Grm2* deletion mouse line were described previously (Johnson et al., 2020a). The floxed mouse line was derived from the same founder as the *Grm2* deletion mouse line. Floxed mice were genotyped by PCR using the primers shown in **Fig. S1**. Briefly, the intron 2 site was genotyped with F1 (5’-CAGATTCTGCTGGCCCATGA-3’) and R1 (5’TTGGTCTCCATTGGATGCCC-3’) primers. This primer set amplifies a 420bp WT amplicon and a 460bp amplicon from the floxed allele. The intron 3 site was genotyped with F2 (5’-AATAGTGCCAGCCAGTGACC-3’) and R2 (5’-CACCTAAATAGAAGTTCTCCC-3’) primers. This primer set amplifies a 352bp WT amplicon and a 392bp amplicon from the floxed allele. The fidelity of the loxP insertions were confirmed by Sanger sequencing of PCR products.

*Grm2^f/f^* mice were crossed with Emx1-IRES-Cre mice (The Jackson Laboratory, stock no. 005628; Gorski et al., 2002) to conditionally delete mGlu_2_ from telencephalon-lineage neurons including cortical neurons. To generate *Grm2^f/f^;Emx1-IRES-Cre^-/-^* and *Grm2^f/f^;Emx1-IRES-Cre^+/-^* mouse for behavioral testing, male *Grm2^f/f^* mice were crossed with female *Grm2^f/f^;Emx1-IRES-Cre^+/-^* mice. In figure legends, genotypes of *Grm2^f/f^;Emx1-IRES-Cre^-/-^* and *Grm2^f/f^;Emx1-IRES-Cre^+/-^* are shorted to *Grm2^f/f^;Cre^-/-^* and *Grm2^f/f^;Cre^+/-^*. Tissue sampling and genotyping for the presence of Cre recombinase were performed by Charles River Laboratories. Experimenters were blind to genotypes during data collection and preliminary analysis.

### Behavioral testing

#### Apparatus

Operant training was conducted in standard mouse operant conditioning chambers equipped with a rod floor, two retractable levers, a food hopper, and a house light (Med Associates, Fairfax, VT). Infrared beams detected head entries into the food hopper. Operant boxes were enclosed in ventilated sound-attenuating chambers. Operant training protocols were controlled through MED PC V software.

Food restriction and reinforcer habituation: Four to seven days prior to onset of operant behavior training, food restriction was initiated and mice were handled and weighed daily. Body weight was gradually reduced to 85-90% of baseline by feeding ∼1.5-2g of standard chow per mouse daily. One day prior to hopper training, mice underwent home cage exposure to palatable food pellets that were used as reinforcers for operant training (20 mg Dustless Precision Pellets, catalog #F0071, Bio-serv). Mice then underwent a single session of hopper training that consisted of non-contingent delivery of food pellets on an RI60 schedule with 50% probability.

Random ratio (RR) and random interval (RI) operant training: Each self-paced operant training session began with concurrent illumination of the house light, which remained on for the duration of the session, and insertion of a single lever. For RR training, mice first underwent three sessions of continuous reinforcement training (CRF) in which each lever press was reinforced with one food pellet. Mice then received two sessions of RR10 training followed by four sessions of RR20 training. Sessions lasted a maximum of 60 minutes and were terminated immediately when a maximum of 5 (CRF day 1), 15 (CRF day 2), or 30 (all remaining sessions) reinforcers were earned. Additional RR20 retraining sessions (2-3) occurred between devaluation, contingency degradation, and progressive ratio assessments. RI training was conducted similar to RR training. All sessions lasted a maximum of 60 minutes and were immediately terminated after 30 reinforcers had been earned. After 3 initial CRF training sessions, mice underwent 2 days of RI30 training followed by 6-10 days of RI60 training (probability 10%). Additional RI60 retraining sessions (3-4) occurred between contingency degradation, omission, and devaluation tests. In rare cases, levers were baited with a crushed pellet early in the training paradigm to encourage exploration of the lever. After each daily training session, mice were provided access to a 20% sucrose solution in their home cage.

#### Devaluation

Sensitivity to changes in outcome value was probed using a sensory-specific satiety protocol consisting of a valued day and a devalued day. Prior to testing, mice were given 1-hour access to food pellets (devalued day) or 20% sucrose solution (used as a control for satiety on the valued day). Immediately following pre-feeding, mice were placed in the operant chamber and lever pressing was measured for 5 minutes under extinction conditions. The order of valued and devalued sessions was counterbalanced between subjects, and a single RR20 or RI60 retraining session was performed between the two devaluation testing days.

Contingency degradation: Sensitivity to changes in action-outcome contingency was evaluated in contingency degradation session (2-3 consecutive sessions). During contingency degradation sessions, conditions were matched to RR or RI training sessions, but reinforcer delivery was no longer contingent on lever pressing. Instead, food pellets were delivered on a fixed interval schedule set to match each subject’s individual rate of reinforcer delivery during the RR20 or RI60 session immediately preceding the first contingency degradation testing. Contingency degradation was performed for 2-3 consecutive sessions.

#### Omission

As an additional measure of sensitivity to changes in action-outcome contingency after RI training, mice underwent omission testing during which lever press responses had to be withheld for 20 seconds to earn a reinforcer. All other conditions were identical to RI training sessions. For **Fig. 3**, all mice underwent 60-minute omission tests. For **Fig. S1**, omission sessions lasted a maximum of 60 minutes or were terminated immediately after 50 reinforcers were earned.

#### Progressive ratio (PR)

As a measure of motivation to obtain the reinforcer, mice underwent a single PR session in which the lever press requirement progressively increased for each reinforcer earned (i.e., 10 lever presses required for the first reinforcer, then 20, 25, 32, 40 etc. following a geometric progression (Richardson and Roberts, 1996). PR sessions lasted a maximum of 6 hours and were immediately terminated if 1 hour lapsed since the last reinforcer was earned. The total number of lever presses and the breakpoint (i.e., the number of lever presses made for the last reinforcer obtained) were recorded.

Deterministic reversal learning in a two-lever place discrimination task: Following hopper training, mice underwent 3 self-paced training sessions that begin with illumination of the house light and insertion of 2 levers, both of which were active. Mice earned a food reinforcer for a single press on either lever (FR1 training). The preferred lever during FR1 training was determined for each mouse, and mice were assigned the non-preferred lever as the active lever for subsequent acquisition of trial-based responding. During the acquisition phase, daily training sessions consisted of 30 trials that began with illumination of the house light followed by insertion of the active and inactive levers after 0.5 seconds. One press of the active lever resulted in reinforcer delivery and ended the trial (retraction of levers and extinguishing of the house light). A press on the inactive lever terminated the trial without reinforcer delivery. If 10 seconds elapsed without a response, the trial was terminated and recorded as an omission. All trials had a 20-second inter-trial interval. After 12 acquisition training sessions, the active and inactive levers were reversed. The reversal phase consisted of 15 sessions with 20 trials per session.

Assessment of locomotor activity in an open field: Mice were placed into a novel gray-walled open field apparatus (40 cm x 40 cm x 35 cm; Stoelting). Ambient light was 15-20 lux. Activity was recorded for 30 minutes by an overhead camera. Mice used for locomotor activity assessment had previously undergone operant training and were returned to *ad libitum* feeding prior to activity assessment.

### Drugs

LY379268 was obtained from Tocris Bioscience (Minneapolis, MN). Stocks were prepared at 10 mM in 0.01N NaOH and frozen at -20°C until use. For intraperitoneal injections, LY379268 was diluted to 0.1 mg/mL in sterile water and administered in a volume of 10 mL/kg. Injections were given 30 minutes prior to behavioral testing. For the experiment assessing LY379268 effects on responding after outcome devaluation, mice were given one hour of pre-feeding time prior to the injection, then an additional 30 minutes of pre-feeding time following the injection. Mice were habituated to vehicle injections during the acquisition phase of the experiment.

### Measurement of *Grm2* expression levels by quantitative polymerase chain reaction (qPCR)

Mice that had previously undergone behavioral testing were anesthetized with isoflurane and decapitated, and brains were removed and rinsed in cold 0.9% NaCl. One-mm coronal sections were prepared using a stainless steel brain matrix, and samples from motor cortex and thalamus were placed in RNAlater (Invitrogen), stored at 4⁰C for 48 hours, then removed from RNAlater and frozen at -20⁰C prior to RNA extraction. RNA was extracted using the RNeasy Lipid Tissue Mini Kit (Qiagen), and RNA concentration was measured using a NanoDrop One (Thermo Fisher Scientific). RNA (400 ng) was reverse transcribed using the Quantitect Reverse Transcription Kit (Qiagen). qPCR reactions contained 1.2 µl cDNA template, 1 µl TaqMan Gene Expression Assay (Mm01235832; Thermo Fisher Scientific), 10 µl TaqMan Fast Advanced Master Mix (Thermo Fisher Scientific), and 7.8 µl Ultrapure water (Invitrogen). qPCR was run on a CFX96 (BioRad) using the following settings: 50°C for 2 minutes, 95°C for 2 minutes, and then 40 cycles of melting at 95°C for 1 second and annealing/extension at 60°C for 40 seconds. Reactions were run in triplicate for 7-12 animals per group. Analysis was performed using the 2^−ΔCT^ method (Schmittgen and Livak, 2008). The CT value was determined by identifying the cycle number at which the fluorescence increased above background. Relative *Grm2* expression was quantified using Actb as an internal control (Applied Biosystems #4352341E). Thus, ΔCT = CT (*Grm2*) − CT (*Actb*).

### Data analysis and statistics

Analysis of operant behavior data was performed using Microsoft Excel and custom Python programs (code available upon request). Statistical tests and data visualization were performed using GraphPad Prism 9.4. For individual mice, lever press bouts during RR or RI responding were defined as a series of 3 or more lever presses with inter-press intervals less than the average inter-press interval across the full session (Renteria et al., 2021). Lever presses during outcome devaluation were normalized for each subject by dividing the lever presses for the valued or devalued session by the total number of presses across both sessions. For contingency degradation tests, lever press rates for each subject were averaged across sessions. Distance travelled and time spent in the center 20 cm x 20 cm portion of the open field apparatus were analyzed using Topscan software (CleverSys). *Grm2* expression was compared between genotypes using an unpaired t test. Because we routinely observe substantial sex differences in operant response rates between wild-type C57BL/6J male and female mice, for each type of operant experiment, we first ran a 3-way ANOVA on responses during training (factors were genotype [between subject], sex [between subject], and session [within subject]) to confirm that there was a main effect of sex, then used 2-way repeated measures (RM) ANOVA (with *post-hoc* Sidak’s multiple comparison tests when appropriate) or unpaired t tests to separately probe the effect of *Grm2* deletion on operant responding and tests of behavioral flexibility in male and female mice. Measurements obtained in open field testing were analyzed by 2-way ANOVA with factors sex and genotype (both between subject). Geisser-Greenhouse corrections were applied to repeated measures analysis when sphericity was not met. For all experiments, n represents the number of mice. Effects were considered statistically significant at p < 0.05. For clarity, only statistical results that are directly relevant to the major questions being asked are presented in the text.

## Acknowledgements

This work was supported by U.S. National Institutes of Health grants AA025403 (K.A.J.), AA010422 (G.E.H.), and AA020889 (G.E.H.). We thank Dr. Angela Thomas and the animal care technicians at Charles River Laboratories for their excellent support of our breeding colony. We also thank the USU Preclinical Behavior and Modeling Core, particularly Dr. Mumeko Tsuda and Ms. Laura Tucker, for access to resources necessary for these experiments. We thank Carolyn Ferguson (University of Pittsburgh) for expert technical support and Dr. David Kupferschmidt (National Institute of Neurological Disorders and Stroke, US National Institutes of Health) for comments on the manuscript.

