## Supplemental Figures for "Assessment of behavioral flexibility in mice with conditional deletion of metabotropic glutamate receptor 2 from *Emx1*-lineage neurons"

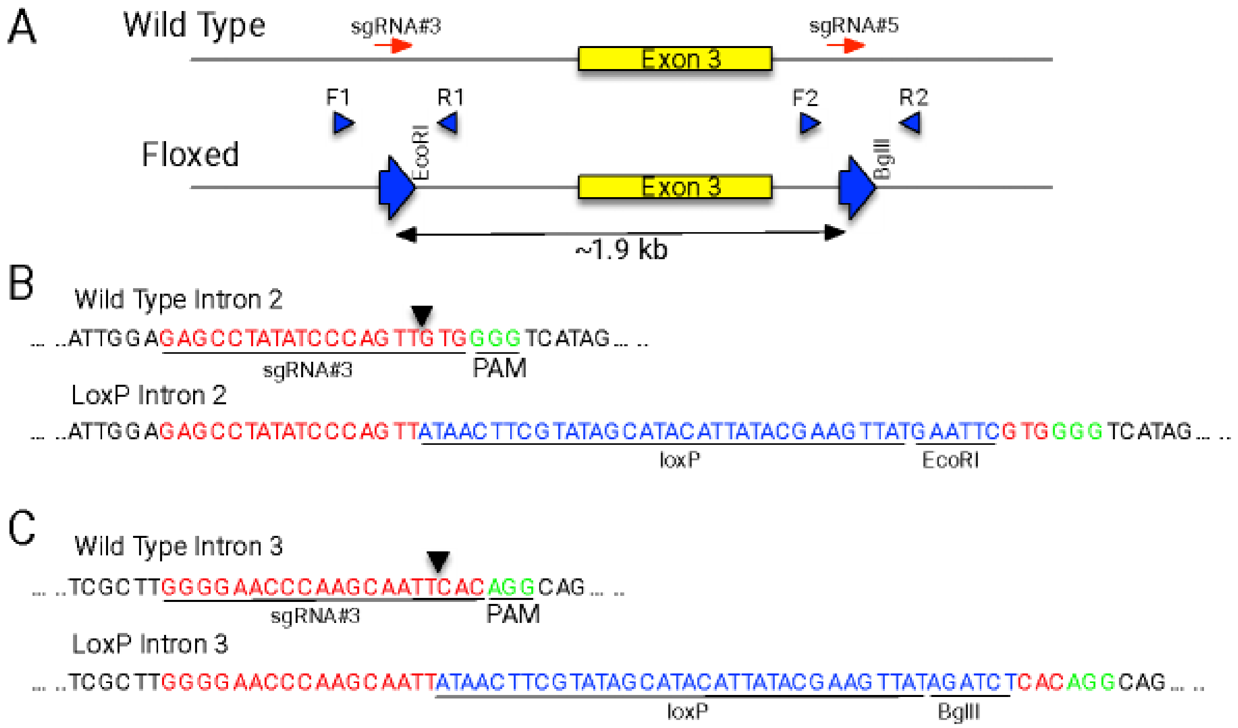


**Figure S1.** Diagram of floxed *Grm2*. (A) Overview of wild-type and floxed *Grm2* alleles. Two sgRNAs (red arrows) were used to direct Cas9 to introns 2 and 3 and insert loxP sequences (large blue arrows) and restriction enzyme sites (EcoRI, BglII). Also shown are primers for PCR genotype analysis (blue arrowheads). (B) DNA sequence of sgRNA binding sites (red), protospacer adjacent motifs (green), and flanking DNA (black) for wild-type and loxP intron 2 sites. (C) DNA sequence of sgRNA binding sites (red), protospacer adjacent motifs (green), and flanking DNA (black) for wild-type and loxP intron 3 sites.

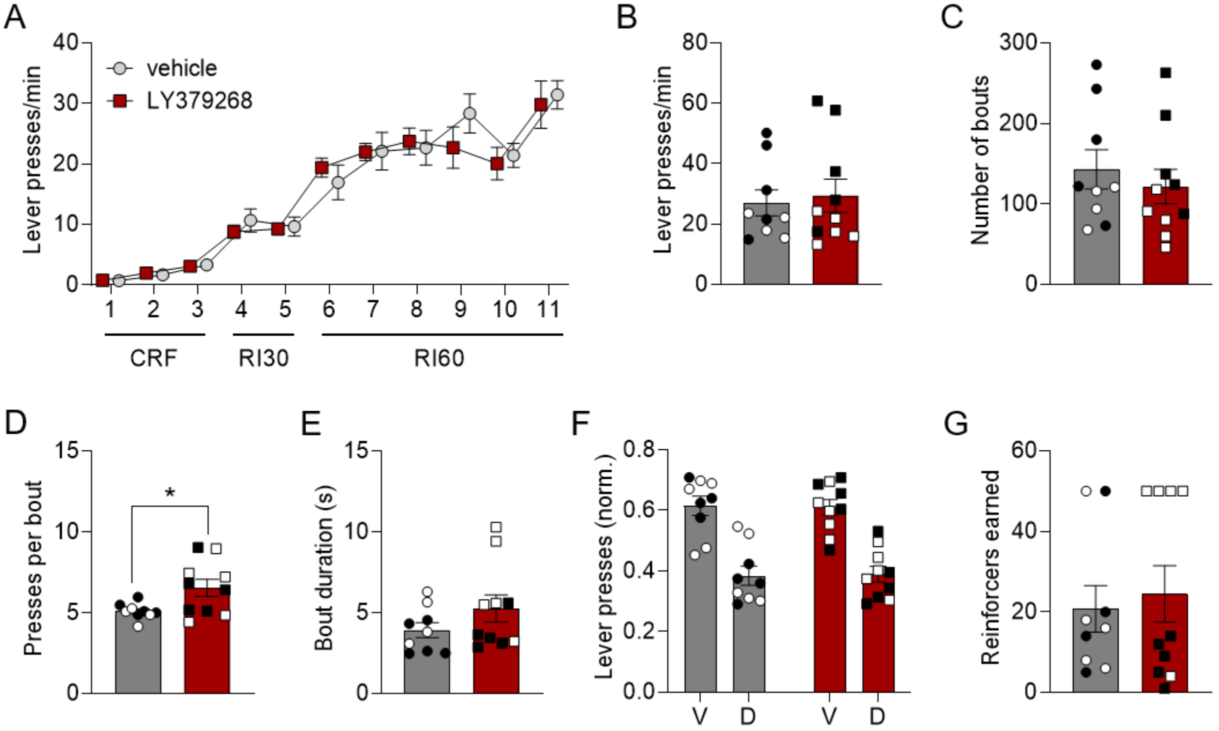


**Figure S2. Evaluation of $mGlu_{2/3}$ agonist effects on RI60 responding, outcome devaluation, and omission performance.** (A) Lever press rates during acquisition of lever pressing on a self-paced random interval schedule. No drugs were administered during the acquisition phase. n = 9 (5 male, 4 female) for the saline group and 10 (5 male, 5 female) for the LY379268 group. (B-E) Lever pressing during an RI60 session following vehicle or LY379268 (1 mg/kg) administration. Measures include press rates across the whole session (B), the number of bouts per session (C), the average number of presses in each bout (D), and the average duration of each bout (E). *$p < 0.05$, unpaired t test. (F) Number of lever presses during valued (V) and devalued (D) sessions following vehicle or LY372968 administration. (G) Number of reinforcers earned during the omission probe following vehicle or LY379268 administration. For this experiment, reinforcers were capped at 50. Data are presented as mean ± SEM with data points from individual mice overlaid for all bar graphs. Data points from male mice are shown as black circles (vehicle) or black squares (LY372968). Data points from female mice are shown as white circles (vehicle) or white squares (LY379268).
